# Dissecting Phylogenetic Support: Unified Decay Indices, AU Tests, and Branch-Site Specific Visualizations

**DOI:** 10.64898/2025.12.05.692543

**Authors:** James O. McInerney, Christopher J. Creevey, Mary J. O’Connell

## Abstract

Phylogenetic inference relies on robust measures of branch support to assess the reliability of evolutionary relationships. While bootstrap resampling and Bayesian probabilities have become the predominant support metrics in molecular phylogenetics, decay indices (also known as Bremer support) provide an alternative perspective by quantifying the amount of evidence required to collapse specific clades. We present panDecay, a unified software framework that extends the classical parsimony-based decay index concept to maximum likelihood (ML) and Bayesian inference approaches. panDecay automates the calculation of ML-based decay indices using log-likelihood differences, performs Approximately Unbiased (AU) statistical tests, and computes Bayesian decay indices through marginal likelihood comparisons using stepping-stone sampling. The software integrates with established phylogenetic tools (PAUP* and MrBayes) and provides comprehensive output including site-specific support analysis, multiple annotated tree formats, and detailed statistical reports. panDecay supports diverse data types (DNA, protein, and discrete morphological characters), offers flexible model specification, and provides parallel processing capabilities for computationally intensive Bayesian analyses. By unifying decay index calculations across parsimony, likelihood, and Bayesian frameworks within a single command-line interface, panDecay enables researchers to obtain complementary support metrics that provide deeper insights into phylogenetic signal strength and distribution across sequence alignments. The software is implemented in Python 3.8+, distributed via PyPI, and released under the MIT license.

## Introduction

Phylogenetic studies are fundamental to understanding organismal evolutionary history as well as the history of genes, and genomes (Felsenstein 2003) and they underpin several areas of biological research, from predicting gene function (Barker and Pagel 2005) to informing drug development and forensic analyses (Hillis 1997). Popular phylogeny inference methods include maximum parsimony (MP), distance matrix (DM), maximum likelihood (ML), and Bayesian inference (BI). These methods are based on explicit mathematical descriptions of how character states transform over time (Semple and Steel 2003). The arrival of high-throughput sequencing technologies has revolutionised phylogenetics, leading to an exponential increase in data volume. Research has shifted from single-gene analyses to phylogenomics (Eisen and Fraser 2003; Burki 2014), where datasets often comprise hundreds, or thousands of taxa and millions of nucleotides or amino acid sites. This genome-scale data initially promised to be a substantial advance, seemingly eliminating data availability concerns and stochastic error as limiting factors for resolving deep phylogenetic questions (Dunn, et al. 2008). However, this expansion in data quantity has simultaneously amplified challenges related to systematic error (Felsenstein 1978) and model misspecification (Jermiin, et al. 2020; Magee, et al. 2021), and substantially increased computational complexity. This creates a scenario where more data, while offering unprecedented resolution, also exposes deeper, often unaddressed, methodological flaws and biological complexities (Magee, et al. 2021), implying that simply accumulating more data is insufficient for achieving accurate phylogenetic inference. Instead, improvements in sophisticated evolutionary models and computationally efficient algorithms, as well as investment in understanding supports and conflicts in phylogenetic data are equally, if not more, critical. Advances in sequence evolution modelling now account for complex evolutionary processes, such as non-stationary, non-reversible, non-homogeneous conditions (Foster 2004), and gene tree-species tree discordance (Larget, et al. 2010), which become more apparent with larger datasets.

Phylogenetic trees are the primary visual representation of inferred evolutionary relationships and are routinely annotated with branch support values (Felsenstein 2003). These values are typically used for quantifying the uncertainty associated with specific phylogenetic hypotheses and for assessing the reliability of the inferred tree topology. The bootstrap approach of sampling characters in a data matrix, with replacement, remains the most widely used measure of branch support in phylogenetic tree reconstruction (Felsenstein 1985). Despite difficulties concerning its statistical interpretation and known limitations, the original bootstrap method continues to be extensively used. A common empirical guideline, suggested by Hillis and Bull (1993), is to use a 70% bootstrap support threshold as an indication of reliability. However, this threshold was established under very specific, perhaps idealised, conditions, such as equal rates of change and symmetric phylogenies.

A significant limitation of Felsenstein’s bootstrap, particularly with the increasing number of taxa in modern datasets, is its tendency to yield very low support values for deep branches (Efron, et al. 1996). This issue stems from the method’s core methodology: a replicated branch must match a reference branch *exactly* to be counted towards its support value. The presence of “rogue” taxa (Russo and Selvatti 2018) with unstable phylogenetic positions perhaps because of deep divergence times, or biological factors like convergence, recombination, or even data errors - can severely depress bootstrap support values. Even if a branch is largely consistent across bootstrap replicates, the misplacement of just one or a few rogue taxa can lead to drastically reduced bootstrap support (Smith 2021). Furthermore, the bootstrap method is computationally intensive, typically requiring analysis times 100-1,000 times longer than phylogenetic inference from the original dataset. While parallelisation and the development of faster algorithms have helped mitigate this problem, it remains a substantial computational burden for the massive datasets common in phylogenomics.

The statistical meaning of traditional Felsenstein’s bootstrap proportions (FBPs) is a subject of intense debate, often interpreted as a measure of repeatability rather than a true confidence level (Soltis and Soltis 2003). A significant issue arises with large datasets where FBPs tend to yield very low supports, even for branches that are highly likely to be correct (Lemoine, et al. 2018). This suggests that the conventional interpretation and utility of support values, particularly bootstrap proportions, are becoming problematic in the phylogenomic era. This observation necessitates a fundamental re-evaluation of how researchers interpret and communicate phylogenetic confidence. Relying solely on traditional bootstrap values, especially in the context of large datasets, can lead to a significant underestimation, or overestimation, of true support or a misinterpretation of underlying phylogenetic signals. Here, we suggest that it is of value to adopt and report a plurality of metrics, in the expectation that this plurality will reflect the statistical certainty, or the strength of evolutionary evidence for a given branch more completely.

Novel approaches are continually being developed to address the limitations of traditional branch support methods. Transfer Bootstrap Expectation (TBE), for example, is a more recent innovation in phylogenetic bootstrap that measures the presence of inferred branches in replicates using a gradual “transfer” distance, as opposed to the binary presence/absence index of Felsenstein’s bootstrap (Lemoine, et al. 2018). TBE has been shown to yield higher and more informative support values than the standard approach, while maintaining a very low rate of falsely supported branches. It is particularly robust to high taxon sampling, a scenario where FBP is negatively impacted. The Bayesian bootstrap allows branch support to be interpreted directly as a posterior probability, which can be more intuitive (Lemoine and Gascuel 2024). Approximate Likelihood Ratio Tests (aLRT) provide faster alternatives to traditional bootstrap for assessing local branch support (Anisimova and Gascuel 2006). Another method proposed as an alternative to FBP is aBayes support, though it is noted to be difficult to apply to large datasets and sensitive to model misspecification (Anisimova, et al. 2011).

The Approximately Unbiased (AU) test was developed to reduce bias in general hypothesis testing, particularly in the context of maximum-likelihood tree selection (Shimodaira 2002). It employs a multiscale bootstrap technique and aims to provide greater accuracy, and a simpler implementation compared to earlier methods like the Shimodaira-Hasegawa (SH) test (Shimodaira and Hasegawa 2001). A key feature of the AU test is its ability to adjust for selection bias, which commonly arises when comparing multiple trees simultaneously and can lead to overconfidence in incorrect tree topologies. The test establishes a null hypothesis that the distributions of site likelihoods for different candidate trees (or roots) are statistically equivalent, and it returns a p-value for each tested tree or root. The primary application of the AU test is used to distinguish, from a set of user-obtained trees, those phylogenetic hypotheses that are, and those that are not significantly worse reconstructions of the phylogeny than the maximum likelihood tree. While the AU test is designed to reduce bias, its asymptotic theoretical foundation assumes a smooth boundary for hypothesis testing, which may not perfectly hold for the complex polyhedral convex cone formed by tree selection problems (Shimodaira 2002). Empirical studies show that instability in tree topology, even if not always statistically significant by the AU test, occurs frequently when new taxa are added to large datasets (Collienne, et al. 2024).

Bootstrapping is broadly applicable and conceptually simple but computationally demanding and its statistical interpretation is ambiguous. The AU test aims for reduced bias and higher accuracy by employing complex multiscale bootstrap techniques, which also involve significant computational resources. This implies that there is no single “optimal” branch support method universally applicable to all phylogenetic analyses. It is necessary to carefully navigate this trade-off, making informed decisions based on their specific dataset characteristics (size, complexity), available computational resources, and the precise biological questions being addressed.

In the context of large phylogenomic datasets, systematic errors can inflate support values for incorrect topologies, meaning that even high support values may not reflect true accuracy. This suggests that high support values, particularly those approaching 100% in genome-scale datasets, are not always reliable indicators of phylogenetic truth (Simmons and Norton 2014). Instead, they can be artifacts of methodological biases, model misspecification, or the sheer volume of data reducing stochastic error without addressing systematic error. As noted for decades, the emphasis when analysing large datasets must shift from *obtaining* high support (which, depending on the method might simply reflect the fact that the dataset is huge in size) to *critically interpreting* its meaning within the broader context of the data and the analytical methods employed. These ongoing challenges have motivated numerous attempts to disassemble phylogenetic signals and noise as well as addressing artefacts, biases and aggregate support measures, for example (Bandelt and Dress 1992; Charleston 1998; Wilkinson 1998; McInerney and Wilkinson 2005; Cummins and McInerney 2011).

The parsimony-based Decay Index, widely known as Bremer support (Bremer 1994) or sometimes referred to as patristic difference (PD) support metric (Grant and Kluge 2008), is a method for quantifying branch support in parsimony-based phylogenetic analyses. It measures the robustness of a clade by calculating the difference in tree length (*i.e*., the number of evolutionary steps) between the most parsimonious tree (MPT) that *lacks* that specific clade and the overall MPT. Essentially, it quantifies how many additional evolutionary steps are required to collapse a particular branch in the tree, or to support the best-fitting alternative topology that contradicts that branch. A higher decay index value indicates stronger support for the branch, as it implies a greater cost (more steps) for its removal or contradiction.

Lee and Hugall proposed a Bremer support-like approach for maximum likelihood (Lee and Hugall 2003). However, their approach related to partitioned datasets. In the more straight-forward context, likelihood support for a particular clade is quantified as the difference in log-likelihood scores between the optimal tree topology that *includes* that clade and the optimal topology that *excludes* it. This provides a likelihood-based equivalent to the parsimony “steps” difference. An equivalent support measure can be obtained in a Bayesian inference context, where the *average* probability from the set of best trees is compared, using the stepping stone approach (Xie, et al. 2011), to the set of trees that do not include the relationship being tested. Therefore, theoretically sound methods exist, but their detailed, standardised implementation in widely accessible software is lacking. Practical barriers of this nature often hinder the widespread adoption of potentially valuable support metrics, despite their theoretical appeal.

To address the practical gap between theoretical frameworks and accessible implementations in phylogenetic support assessment, we have developed PanDecay, a Python command-line tool that implements parsimony, maximum likelihood and Bayesian based decay indices for phylogenetic branch support assessment. The software extends the classical parsimony-based Bremer support concept to the ML and Bayesian frameworks by automating the calculation of differences between optimal ML trees and sets of best Bayesian trees, with their constrained alternatives where specific clades are forced to be non-monophyletic. PanDecay is available under an MIT Open Access licence from https://github.com/mol-evol/panDecay/.

In line with the philosophy of deconstructing phylogenetic signals, biases and methodological approaches, PanDecay integrates multiple analytical methods, enabling systematic evaluation of tree topology confidence across all nodes in a phylogeny. The software leverages PAUP* (Swofford 2003) as its primary computational engine for ML and parsimony analyses, while integrating MrBayes (Ronquist, et al. 2012) for Bayesian analysis.

The program operates through an integrated five-phase analysis pipeline: (1) optimal tree reconstruction under the specified optimality criterion with likelihood scoring, (2) systematic generation of topological constraints for each internal branch using PAUP*’s converse=yes functionality to force non-monophyly of target clades, (3) constrained tree searches under identical model parameters to identify the best trees that violate each monophyletic grouping, (4) calculation of support metrics including log-likelihood differences (ML decay), marginal likelihood differences (Bayesian decay), and parsimony step differences, and (5) statistical hypothesis testing using the Approximately Unbiased (AU) test for ML analysis, Bayes decay for Bayesian analysis, and traditional parsimony support thresholds, as well as Felsenstein bootstrapping.

PanDecay’s implementation of a novel Bayesian approach to branch support assessment proceeds through what we term “Bayesian Decay” (BD) indices. This method extends the decay index concept to Bayesian phylogenetic inference by quantifying the difference in marginal likelihoods between models that enforce and prohibit specific clades. The Bayesian decay analysis operates through the following process: (1) an unconstrained Bayesian MCMC analysis is performed to estimate the marginal likelihood of the data under the specified model, (2) for each internal branch obtained from the original maximum likelihood tree, a negative constraint is applied that prohibits the taxa in that clade from forming a monophyletic group, (3) a constrained MCMC analysis is run to estimate the marginal likelihood under this restriction, (4) the Bayesian Decay value is calculated as the difference in log marginal likelihoods (unconstrained minus constrained), and (5) Bayes Factors are computed as the exponential of the BD values.

PanDecay employs stepping-stone sampling (Xie, et al. 2011) as the default method for marginal likelihood estimation, which has been shown to provide more accurate estimates than the harmonic mean estimator. The software interfaces with MrBayes (Huelsenbeck and Ronquist 2001; Ronquist, et al. 2012) to perform the MCMC analyses, supporting complex evolutionary models including GTR+G+I for nucleotide data and various amino acid substitution models for protein sequences.

PanDecay provides comprehensive support for diverse data types including DNA sequences, protein sequences, and discrete morphological characters. The software implements flexible model specification with support for standard DNA models (*e.g.* GTR, HKY, JC), protein models (*e.g.* JTT, WAG, LG), and discrete character models (Mk), all with optional extensions to include gamma-distributed rate heterogeneity (+G) and proportion of invariable sites (+I). Model parameters can be estimated or fixed, with support for empirical base frequencies and custom PAUP* command blocks for complex model specifications. The tool integrates bootstrap analysis alongside decay calculations, enabling researchers to compare these fundamentally different approaches to branch support assessment within a single analytical framework.

A distinctive feature of PanDecay is its comprehensive site-specific likelihood analysis capability. When enabled through the `--site-analysis` option, the software performs character-level decomposition of phylogenetic signal by calculating site-by-site likelihood differences between optimal and constrained trees for each tested branch. This analysis generates detailed output including supporting versus conflicting site counts, support ratios, weighted support ratios based on the magnitude of likelihood differences, and sophisticated visualisations showing the distribution of site-specific evidence across the alignment. The tool creates individual site analysis reports for each branch, histograms showing the distribution of site-specific likelihood differences, scatter plots revealing patterns of support across alignment positions, and summary tables quantifying the overall distribution of phylogenetic signal.

PanDecay implements a multi-modal analysis system that enables simultaneous evaluation of branch support under different optimality criteria. The software can perform coordinated ML, Bayesian, and parsimony analyses `(--analysis all`), recognising that optimal trees may differ between criteria while applying identical topological constraints across all analyses. This approach provides researchers with a comprehensive view of branch support that reveals both convergent and divergent signals across different phylogenetic frameworks.

PanDecay generates an extensive suite of output files designed for both human interpretation and programmatic processing. The primary output includes easily interpreted, formatted Unicode tables, with standardised significance indicators, comprehensive markdown reports with analysis parameters and interpretation guidelines, and multiple annotated tree formats compatible with visualisation software such as FigTree (https://github.com/rambaut/figtree/). PanDecay creates five distinct tree annotation types: AU p-value trees, log-likelihood difference trees, combined annotation trees, bootstrap support trees, and comprehensive trees integrating all support measures. Site-specific analysis results are presented through detailed tabular summaries, individual branch reports, and integrated matplotlib visualisations including histograms and scatter plots.

ML decay indices are calculated by subtracting the constrained tree log-likelihood from the ML tree log-likelihood. Consequently, larger positive values indicate stronger support for the original clade. Statistical significance is assessed separately through AU tests, with p-values testing the null hypothesis that there is no statistically significant difference between the ML tree and the constrained alternatives. Bayesian decay analysis employs marginal likelihood estimation through the stepping-stone sampling (Xie, et al. 2011) approach, with comprehensive convergence diagnostics including ESS, PSRF, and ASDSF monitoring (Ronquist, et al. 2012). PanDecay provides detailed interpretation guidelines emphasising that support values should be evaluated within the context of model adequacy and potential systematic errors.

The software requires Python 3.8+ with standard scientific libraries (BioPython, NumPy, matplotlib, seaborn for visualisations) and functional installations of PAUP* and MrBayes. It is compatible with Unix-like operating systems (Linux, macOS) and includes comprehensive error handling, timeout protection, and debugging capabilities. The tool supports parallel processing through multiple threads for PAUP* (Swofford 2003) operations and MPI support for MrBayes analyses, and using the Beagle library (Ayres, et al. 2012), can move the calculations onto GPU processors, making it suitable for large-scale phylogenomic studies.

Figure 1 presents the results of a basic analysis of primate mitochondrial DNA phylogeny (Hayasaka, et al. 1988) where we can explore of the relationship between Bootstrap Support (BS), Δlog-likelihood (ΔLnl), branch length (Brlen) and AU-Decay probability (AU), Bayes-Decay (BD) and Parsimony-Decay (PD) analyses. This exemplar analysis was not designed to identify the best-fitting model of nucleotide substitution; instead, we used the Jukes-Cantor (1969) model for the sole purpose of illustrating how various support measures could interact. Internal branch lengths are shown on the tree, rounded to four decimal places.

**Figure 1:**
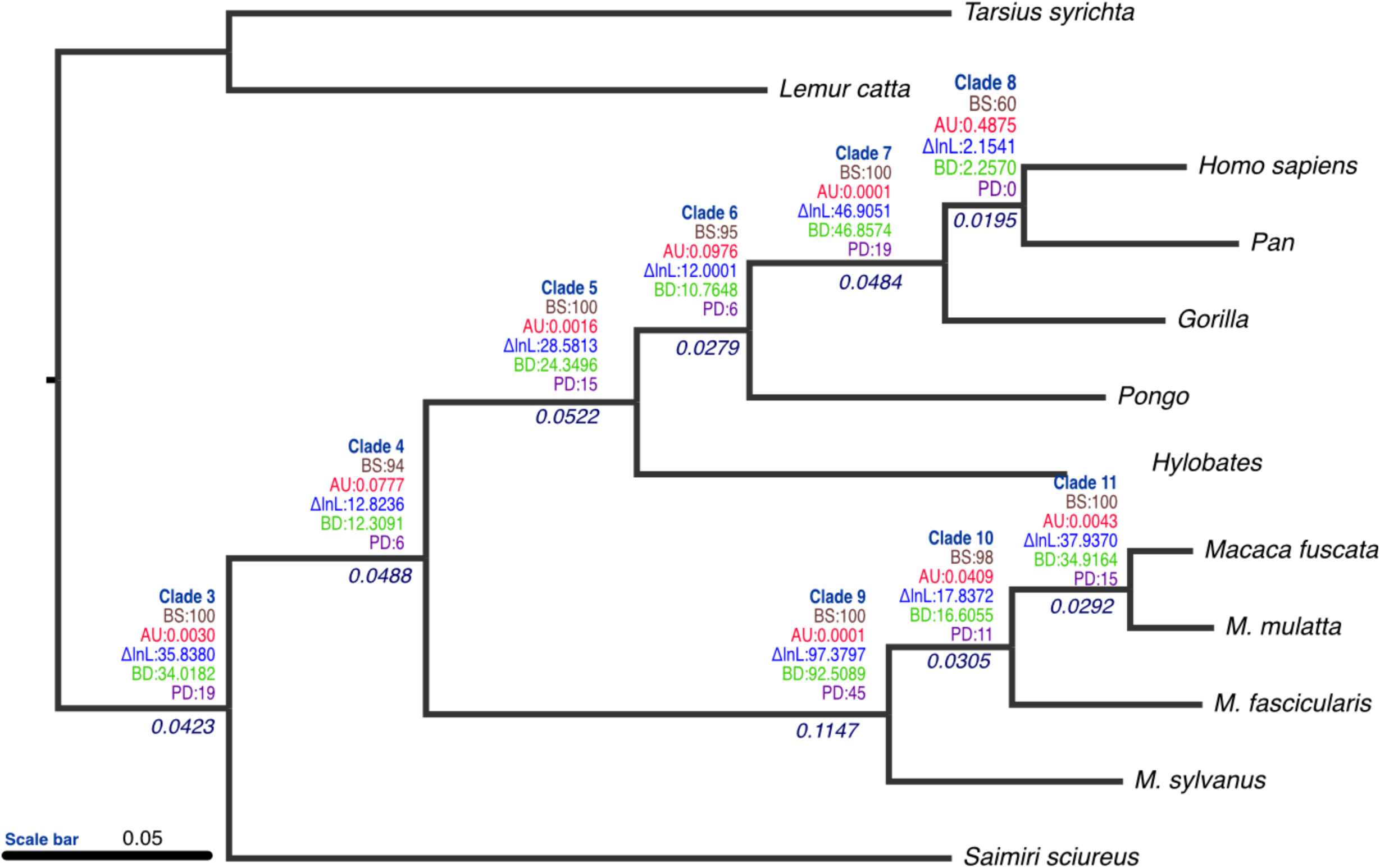
Primate phylogeny inferred using Maximum Likelihood, with internal branch lengths indicated under the relevant branch, while PanDecay clade numbers, Bootstrap Support (BS) values, AU test result (AU), log-likelihood difference (Λ1lnL), Bayesiand Decay (BD) and Parsimony Decay (PD) values indicated above the relevant branch.

Starting with the clade that groups humans and chimpanzees (clade 8 in figure 1), the internal branch length is short - just 0.0195 substitutions per site. This clade has a Δlog-likelihood score of 2.1541, receives weak bootstrap support (60%) that would likely be disregarded by most phylogenecists, a Bayesian decay of 2.2570, a Parsimony decay value of zero, and is not supported by the approximately unbiased (AU) test (P = 0.4875). In contrast, the next internal branch - uniting humans, chimpanzees, and gorillas (labelled clade 7 in figure 1) - tells a different story. The branch length is more than twice as long, and the clade receives 100% bootstrap support, while the AU test strongly rejects the best tree that doesn’t contain this branch (P ≈ 0), Δlog-likelihood score is much higher at 46.9, the Bayes-Decay value is 46.85, while the parsimony decay value is 19. If we were comfortable with the model of sequence evolution used to evaluate this tree, then we would have several consilient lines of evidence that would support the retention of this branch on the tree.

The clade including the orangutan (clade 6 in figure 1) has a shorter internal branch (0.0279 substitutions per site), but received 95% bootstrap support, which many would consider to be strong support for this grouping. Despite receiving 95% bootstrap support, it is not supported by the AU test (p = 0.0976). This illustrates an important case where bootstrap support appears strong, yet the AU test is not in agreement, on the same dataset and model. The Δlog-likelihood score is 12.0, and the bayes decay value is 10.76. The values overall for clade 6 and the deeper clade, clade 4 seem to be somewhat in agreement for most of these indices (both have strong bootstrap support, both fail the AU test), but there is a considerable difference in branch lengths subtending these two clades – clade 6 has a branch length of 0.0279 subs/site, while clade 4 0.0488, almost twice as long. The branch length for clade 4 is very similar to the branch length for clade 3 (0.0488 versus 0.0423), but clade 3, with its shorter branch length, passed the AU test, and indeed, by all measures has more statistical support.

These examples highlight that while bootstrap support, Δlog-likelihood, parsimony and bayes decay scores, branch lengths, and AU test results are all correlated in many ways, no single measure gives a complete picture. Instead, a full understanding emerges only when all metrics are considered together.

The plot in figure 2 presents the results of a site-specific likelihood difference analysis for a particular clade in the phylogenetic tree - clade 11, which groups *Macaca fuscata* with *M. mulatta*. Each bar in the figure represents the difference in log-likelihood for a given site when comparing the optimal (unconstrained) tree to a tree in which this clade is constrained. Green bars indicate that a site supports the unconstrained tree more strongly; the bar’s length reflects how much better the site fits the optimal tree. Red bars show sites that favour the constrained tree, meaning their likelihood scores are higher on the best tree where clade 11 is forced to be absent. This type of analysis could be useful for detecting recombination or horizontal gene transfer. For instance, a cluster of consecutive red bars may suggest a region of the alignment that evolved under a different tree topology, consistent with such events. In addition, the PanDecay program will report the relative supports for this clade by reporting the number of sites favouring the unconstrained clade (752 in this analysis), the number that support the best tree that doesn’t contain the clade (146 in this case), the raw ratio of support (5.15) and a ratio of support and conflict that is weighted by likelihood scores (1.87 in this case).

**Figure 2:**
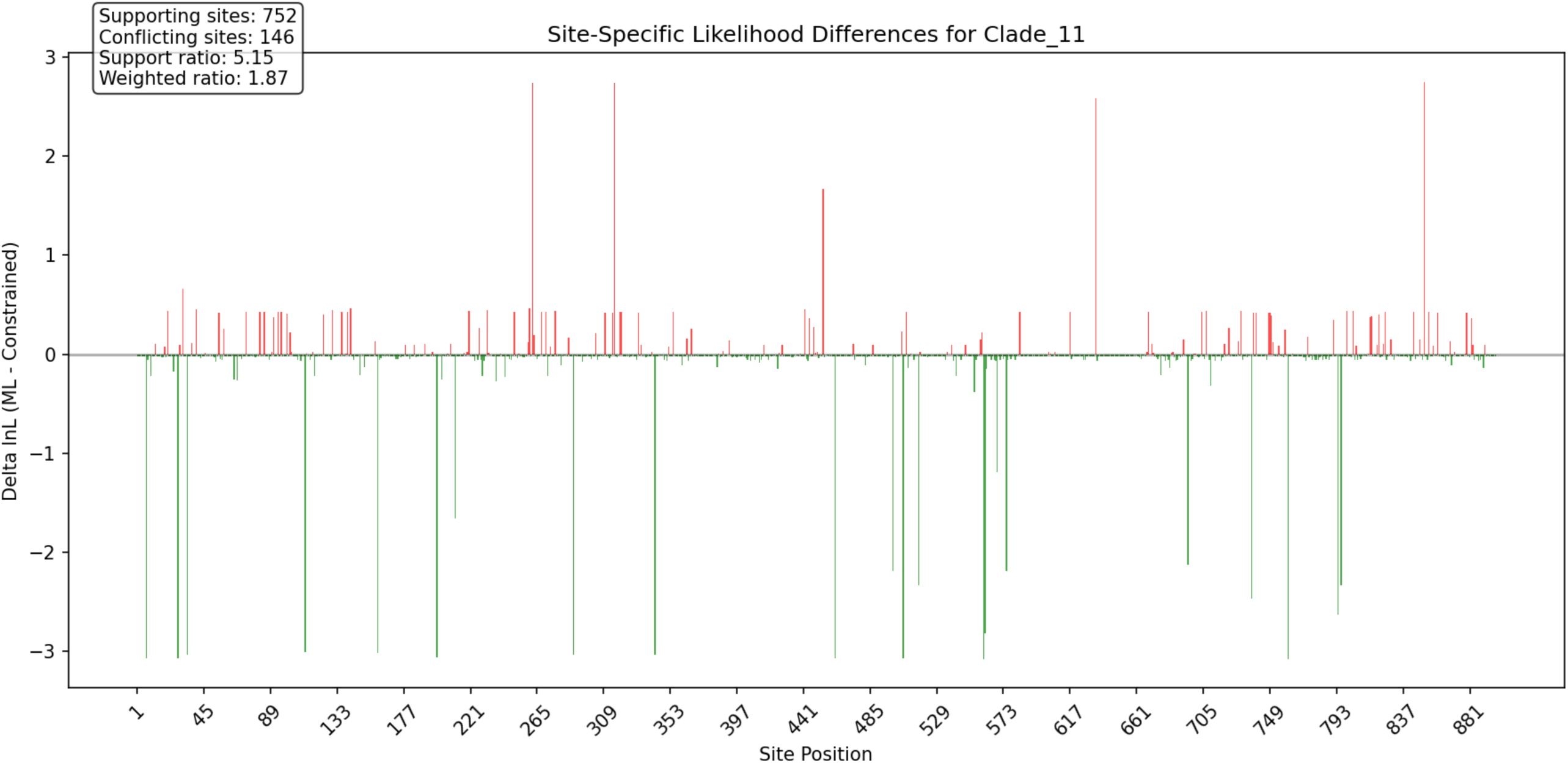
Site-specific, clade-specific analysis of support and conflict for a focal clade. This plot refers to clade 11 on Figure 1. The green bars indicate nucleotide columns where the likelihood is higher for the unconstrained (maximum likelihood) tree, whereas the red bars indicate nucleotide columns that have a higher likelihood for the best tree that doesn’t contain clade 11.

The PanDecay program will output a site-specific figure for every clade on the tree and these plots can be compared to understand whether support comes from different parts of the matrix for different groups on the tree.

The distribution shown in Figure 3, which is derived from Figure 2, indicates that most sites in the data matrix with a likelihood score difference between the constrained and unconstrained trees showed only a small difference. While most sites offered marginal support, a few outliers strongly favoured one tree over the other. As expected, the maximum likelihood (unconstrained) tree received broader overall support. Of the 752 sites that favoured it at this internal node, more than 725 showed only a modest preference - less than 1 log-likelihood unit. However, 17 sites provided strong support, with differences exceeding 3 log-likelihood units in favour of the unconstrained tree. In contrast, the constrained tree was supported by fewer sites - just 146 in total. Although these sites tended to have higher average Δlog-likelihood scores compared to most sites supporting the optimal tree, none exceeded a difference of 3 log-likelihood units. This pattern highlights that strong overall support for the optimal tree is largely due to the cumulative effect of many small contributions, rather than a few highly influential sites. The PanDecay program will produce an output of this kind for every internal branch of the optimal tree.

**Figure 3:**
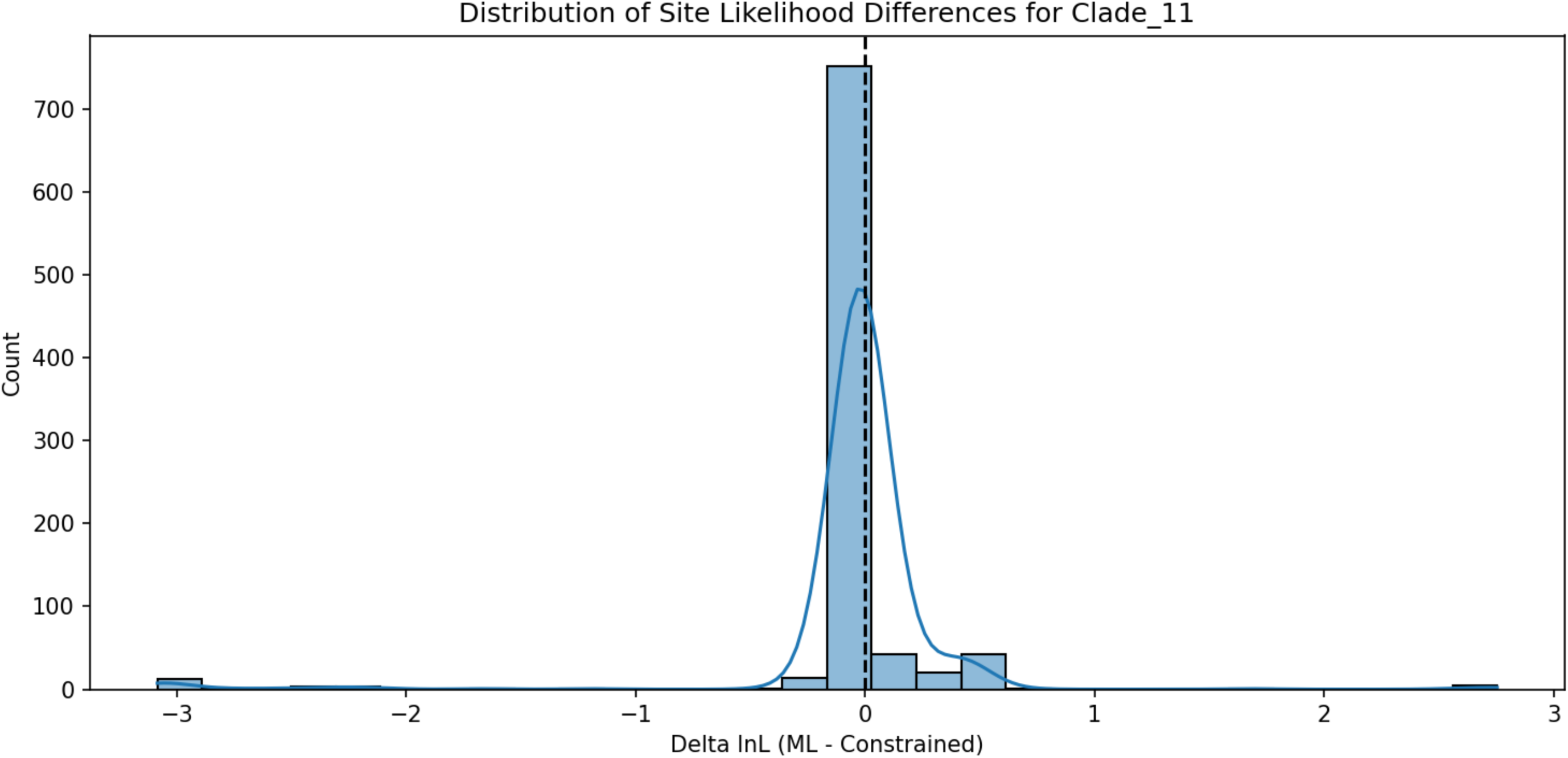
Distribution of site likelihood differences for clade 11. The differences in likelihood score are binned into an appropriate number of bins and the counts are shown on the left.

Summing support and conflict over the tree using maximum likelihood, the panDecay analysis revealed substantial phylogenetic support for the ML tree, across all examined clades, with supporting sites consistently outnumbering conflicting sites by factors ranging from 2.16 to 5.15 (see Table 1). Clade 11 exhibited the strongest support with 752 sites having a higher likelihood on the unconstrained tree versus only 146 sites receiving a higher likelihood score on the constrained tree (support ratio = 5.15), while clade 8 showed the most modest support ratio of 2.24. Neutral sites were rare, appearing in only three clades (3, 5, and 10) with a single site each. Given that the sites are being scored using 64-bit floating point precision, and a threshold value of zero difference, sites must have identical scores on both trees. When incorporating log-likelihood differences between the maximum likelihood tree and best alternative trees, the weighted support ratios ranged from 1.06 (clade 8) to 2.17 (clade 9), indicating that supporting sites generally contributed more strongly to the likelihood than conflicting sites. It is notable that for the human-chimp-gorilla groupings, the number of sites supporting the human-chimp clade was more than twice as many as the number supporting chimp-gorilla, but when weighted by likelihood score, the difference is much closer. This suggests that the majority of “supporting” sites for this clade are providing only a very small amount of support. Clade 11, while having the highest raw number of sites preferring the ML tree, was only in third position when these sites were re-weighted by likelihood score, with clades 9 and 7 being higher by that measure. Clade 3, the deepest split in the tree, had the lowest ratio of raw counts of supporting versus conflicting sites, but when reweighted, it moved to the median position within the nine tested branches.

**Table 1:** Site analysis showing which alignment positions support or conflict with each clade. The raw counts of sites favouring the ML tree versus those favouring the alternative best tree, are given and their support ratio is the ratio of these two numbers. The cumulative negative log-likelihood scores for those sites supporting the ML tree versus those supporting the alternative are used to give a weighted ratio of the support for each clade.

| **Clade ID** | **Supporting Sites** | **Conflicting Sites** | **Neutral Sites** | **Support Ratio** | **Sum Supporting $\Delta$** | **Sum Conflicting $\Delta$** | **Weighted Support Ratio** |
| --- | --- | --- | --- | --- | --- | --- | --- |
| **Clade 3** | 613 | 284 | 1 | 2.1585 | -93.6642 | 57.8261 | 1.6198 |
| **Clade 4** | 631 | 267 | 0 | 2.3633 | -50.8239 | 38.0002 | 1.3375 |
| **Clade 5** | 666 | 232 | 0 | 2.8707 | -70.6060 | 42.0247 | 1.6801 |
| **Clade 6** | 651 | 247 | 0 | 2.6356 | -40.0876 | 28.0874 | 1.4272 |
| **Clade 7** | 693 | 205 | 0 | 3.3805 | -93.4764 | 46.5712 | 2.0072 |
| **Clade 8** | 621 | 277 | 0 | 2.2419 | -39.4304 | 37.2762 | 1.0578 |
| **Clade 9** | 659 | 238 | 1 | 2.7689 | -180.6320 | 83.2521 | 2.1697 |
| **Clade 10** | 664 | 233 | 1 | 2.8498 | -52.0007 | 34.1634 | 1.5221 |
| **Clade 11** | 752 | 146 | 0 | 5.1507 | -81.3266 | 43.3896 | 1.8743 |

## Discussion

The phylogenomic era has challenged traditional approaches to assessing branch support in phylogenetic inference (Collienne, et al. 2024; Lemoine and Gascuel 2024). While the accumulation of vast genomic datasets initially promised to resolve deep evolutionary questions definitively, the reality has been more complex (Dopazo, et al. 2004; Galtier and Daubin 2008; Steenwyk and King 2024). Large datasets can amplify systematic biases and model misspecification, leading to inflated support values that may not reflect true phylogenetic accuracy. This phenomenon necessitates a critical evaluation of how we measure and interpret branch confidence in modern phylogenetic analyses.

PanDecay addresses a gap between theoretical frameworks and practical implementation in phylogenetic support assessment. While Lee and Hugall (2003) established the theoretical foundation for maximum likelihood-based decay indices, no readily accessible software has implemented this approach until now. Our tool provides researchers with a straightforward method to calculate ML decay indices using reverse constraints, offering an alternative perspective on branch support that complements traditional bootstrap, jackknife or Bayesian Inference approaches.

Our analysis of the primate mitochondrial phylogeny demonstrates that different branch support metrics capture distinct aspects of phylogenetic uncertainty. While bootstrap support and AU test p-values showed strong concordance for most branches (r^2^ = 0.87), notable exceptions revealed the limitations of relying on any single metric. For instance, the human-chimp clade received 100% bootstrap support but an AU p-value of 0.086, suggesting that while the data consistently favours this grouping across resampled datasets, alternative topologies cannot be statistically rejected. This discordance illustrates how bootstrap values measure repeatability while AU tests assess statistical distinguishability - fundamentally different concepts that can yield conflicting signals.

The relationship between branch length and support values proved particularly revealing. Short internal branches tended to show lower support across all metrics, confirming that rapid diversification events remain challenging to resolve regardless of the support measure employed. However, the magnitude of this effect varied: bootstrap support showed the steepest decline with decreasing branch length, while decay indices maintained more stable values. This suggests that decay-based metrics may be less susceptible to the branch-length artifacts that plague bootstrap analyses in phylogenomic datasets.

Our implementation of Bayesian decay indices yielded an unexpected but theoretically sound result: BD values closely approximated maximum likelihood differences (ΔlnL). This convergence occurs because both metrics compare models differing only in topological constraints while maintaining identical substitution parameters. The marginal likelihood in such comparisons becomes dominated by the likelihood component, with minimal prior influence. This observation validates the theoretical framework while suggesting that for topology testing, likelihood-based and Bayesian approaches may provide redundant rather than complementary information.

The site-specific analysis revealed substantial heterogeneity in phylogenetic signal across the alignment. While most clades showed support ratios exceeding 2:1, the distribution of supporting sites was highly non-uniform. Clade 11, despite having the highest raw count of supporting sites (752), ranked only third when sites were weighted by their likelihood contributions. This pattern suggests that some branches are supported by many sites with weak signal, while others derive support from fewer but more informative positions. Such insights are impossible to obtain from aggregate support measures alone and highlight the value of character-level signal decomposition.

The practical implications for phylogenomic studies are clear. As datasets grow larger, systematic biases become increasingly problematic, potentially inflating traditional support values for incorrect topologies. Decay indices offer a complementary perspective by quantifying the actual evidence required to contradict a clade rather than merely its repeatability under resampling. The ability to identify which specific alignment positions drive phylogenetic conclusions enables targeted investigation of potentially problematic regions, whether due to alignment errors, recombination, or heterogeneous evolutionary processes.

By decomposing phylogenetic support on a site-by-site basis, panDecay identifies which specific alignment positions contribute to or conflict with each clade’s support. By identifying which alignment positions contribute to, or conflict with, each branch, it may be possible to detect potential recombination events, horizontal gene transfer, or alignment errors that may compromise phylogenetic inference. This granular view of support distribution moves beyond summary statistics to highlight character-level evidence underlying tree topology, enabling more informed interpretation of phylogenetic results.

However, it is important to emphasise that no single support measure provides a complete picture of phylogenetic confidence. The relationship between branch length, likelihood differences, bootstrap support, and AU test results is complex and likely to be context dependent. Some groupings on a tree might have a relatively straightforward relationship between branch length, bootstrap support, Λ1Log-Likelihood and AU test, most likely when there are many supporting sites and few conflicting sites. However, we expect that when support at any node is complicated with a plurality of supporting and conflicting signals, then it is important to tease these apart.

## Acknowledgements

JMc would like to acknowledge the support of The Leverhulme Trust fellowship grant ref RF-2023-408.

## Notes

https://github.com/mol-evol/panDecay

### Competing Interest Statement

The authors have declared no competing interest.

